# Extraction of relations between genes and diseases from text and large-scale data analysis: implications for translational research

**DOI:** 10.1101/007443

**Authors:** Àlex Bravo, Janet Piñero, Núria Queralt, Michael Rautschka, Laura I. Furlong

## Abstract

**Background:** Current biomedical research needs to leverage and exploit the large amount of information reported in publications. Automated text mining approaches, in particular those aimed at finding relationships between entities, are key for identification of actionable knowledge from free text repositories. We present the BeFree system aimed at identifying relationships between biomedical entities with a special focus on genes and their associated diseases.

**Results:** By exploiting morpho-syntactic information of the text BeFree is able to identify gene-disease, drug-disease and drug-target associations with state-of-the-art performance. The application of BeFree to real-case scenarios shows its effectiveness in extracting information relevant for translational research. We show the value of the gene-disease associations extracted by BeFree through a number of analyses and integration with other data sources. BeFree succeeds in identifying genes associated to a major cause of morbidity worldwide, depression, which are not present in other public resources. Moreover, large-scale extraction and analysis of gene-disease associations, and integration with current biomedical knowledge, provided interesting insights on the kind of information that can be found in the literature, and raised challenges regarding data prioritization and curation. We found that only a small proportion of the gene-disease associations discovered by using BeFree is collected in expert-curated databases. Thus, there is a pressing need to find alternative strategies to manual curation to review, prioritize and curate text-mining data and incorporate it into domain-specific databases. We present our strategy for data prioritization and discuss its implications for supporting biomedical research and applications.

**Conclusions:** BeFree is a novel text mining system that performs competitively for the identification of gene-disease, drug-disease and drug-target associations. Our analyses show that mining only a small fraction of MEDLINE results in a large dataset of gene-disease associations, and only a small proportion of this dataset is actually recorded in curated resources, raising several issues on data prioritization and curation. We propose that joint analysis of text mined data with data curated by experts appears as a suitable approach to both assess data quality and highlight novel and interesting information.

## Background

Due to the increasing size of literature repositories, there is a strong need for tools that firstly, identify and gather the relevant information from the literature, and secondly, place it in the context of current biomedical knowledge. Nowadays, the automatic analysis of the literature by text mining approaches eases the access to information otherwise locked in millions of documents and supports translational research projects [1].

Despite of these advances, several challenges remain to be solved in the text mining field, such as the identification of complex relationships between entities of biomedical interest and the exploitation of the extracted information in real-case settings for supporting specific research questions in translational research. This is particularly relevant for researchers interested in human diseases, since they are currently struggled by the large number of publications in their domain. There is a pressing need for methods that can extract this information in a precise manner, can be applied to large document repositories, and are able to provide the extracted data in a way that can be integrated with other pieces of information to aid subsequent analysis and knowledge discovery. In particular, text mining tools that help in the identification of the actionable knowledge from the vast amount of data available in document repositories are key for bridging the gap between bench and bedside [2].

In the past, most efforts in text mining of relationships have been devoted to the identification of interactions between proteins, both due to the availability of corpora and the push driven by specific text mining challenges [1]. In contrast, less attention has been paid to the identification of relationships involving entities of biomedical interest such as diseases, drugs, genes and their sequence variants. In the last years, however, this trend has changed and there is much more interest in gathering this kind of information [3, 4]. There are examples of systems developed for identification of drug-gene interactions [5, 6], drug-drug interactions [7, 8], drug-adverse effect [9, 10], gene-disease [11, 12], and also systems covering different types of relationships [13]. Regarding the specific methodologies available for relation extraction (RE), supervised learning approaches have shown good performance exploiting both syntactic and semantic information [14].

Most of the studies have focused in kernel based methods to identify associations between entities [15–18]. Basically, these methods are able to classify text based on how a relationship between two entities is represented. Different kinds of features, such as word frequencies in the sentences or the relationship between words provided by phrase structure or dependency trees, can be used to represent a relationship between two entities. A common approach involves considering distance criteria like the shortest path between the candidate entities in a parse tree to unravel associations [19, 20].

In this paper, we propose the combination of the Shallow Linguistic Kernel (*K_SL_*) [17] with a new kernel that exploits deep syntactic information, the Dependency Kernel (*K_DEP_*), for the identification of relationships between genes, diseases and drugs. The *K_SL_*, which uses only shallow syntactic information, was successfully applied to extract adverse drug reactions from clinical reports [9] and drug-drug interactions [8]. On the other hand, the *K_DEP_* exploits the syntactic information of the sentence using the walk-weighted subsequence kernels as proposed by [21]. A major requisite of supervised learning approaches for relation extraction is the availability of annotated corpora for both development and evaluation. Although there are several annotated corpora for identification of PPIs (LLL, AIMed, Bioinfer, HPRD50 and IEPA), manually annotated corpora for other associations are scarce [19]. Our group has developed one of these resources, the EU-ADR corpus, that contains annotations on drugs, diseases, genes and proteins and associations between them [22]. We used this corpus to develop a RE system for the identification of relationships between genes and diseases. In addition, we also evaluated the RE systems for the identification of relationships between diseases, drugs and their targets. As we are particularly interested in identifying associations between genes and diseases, we also developed a new corpus in this domain by a semi-automatic annotation procedure based on the Genetic Association Database (GAD), an archive of human genetic association studies of complex diseases and disorders, as a starting point. The RE module in combination with our previously reported Biomedical Named Entity Recognition or BioNER [23] constitutes the BeFree system (http://ibi.imim.es/befree/).

In addition to the evaluation of the performance of the RE system based on Precision (P), Recall (R) and F-score (F), that is common practice in the text mining domain, we wanted to assess the ability of the BeFree system to identify useful information in the context of concrete biomedical problems. More specifically, we applied the BeFree system for the extraction of associations between genes and diseases to two real-case scenarios: 1) the search for genes associated to one of the most prevalent diseases, depression, and 2) the population of DisGeNET, a database of gene-disease associations [24]. In the first case study we demonstrate the ability of BeFree to identify useful information related to this particular disease. In the second case study we evaluated the application of BeFree to large-scale data extraction and integration with another knowledge source. This resulted in a very large dataset on gene-disease associations (approx. 500,000 associations) that raised issues related with the quality of the extracted information. Therefore, we were faced with another challenge, which is the prioritization of the results obtained by large-scale mining of the biomedical literature. Since manual curation is not possible for this kind of large datasets, we performed a series of analysis on the data in order to gain insight on its quality and provide a discussion on their outcomes.

In summary, here we present a novel text mining system, BeFree, specifically focused on the identification of associations between drugs, diseases and genes. Another important contribution of this work is the GAD corpus for the evaluation of RE systems for gene-disease associations. We then focus on the identification of gene-disease relationships, and analyse the outcome of the two case studies that highlight the value of the extracted information, and finally discuss the impact of this kind of approach for translational research. We address some of the current challenges in the field, such as improving relation extraction for entities of biomedical interest, disambiguation between semantically different entities, integration with existing knowledge bases and exploitation of extracted information in real-case scenarios.

The complete set of gene-disease associations extracted by BeFree, with the supporting statements and information on the provenance, is available in DisGeNET (http://www.disgenet.org/). The corpora used in this study, including the new corpus on gene-disease associations, are available at http://ibi.imim.es/befree/#corpora. The BeFree code is available upon request.

## Results and Discussion

We have developed a new RE system to identify associations between genes, drugs and diseases based on the exploitation of semantic and morpho-syntactic information from the text. We first present the results of its evaluation aimed at assessing the performance of both kernels (*K_SL_* and *K_DEP_*, see Figure 1 and 2) using morpho-syntactic features on three relationships, drug-target, gene-disease and drug-disease, using the EU-ADR and GAD corpora (see Table 1 for corpora statistics). The complete set of results is available online at http://ibi.imim.es/befree/#supplbefree. Only a representative set of the results is depicted in the manuscript. We also conducted a series of experiments on the identification of protein interactions in order to evaluate the performance of the *K_DEP_* kernel using different features and compare it with previous results (Suppl. File S1).

**Figure 1.**
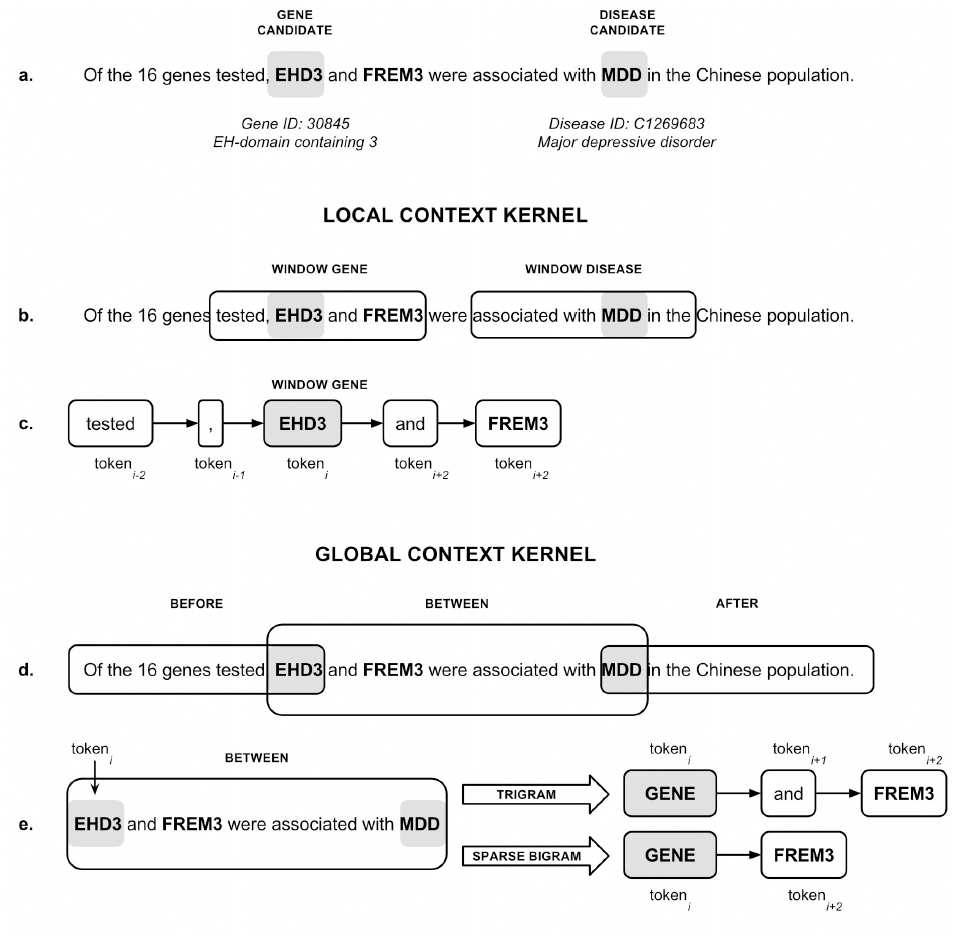
Global and local context kernels to represent a gene-disease association. a) The sentence extracted form a MEDLINE abstract (PMID:22337703) expresses the association between the disease MMD (Major Depressive Disorder) and the genes EHD3 and FREM3. We will focus in the association between EHD3 and MMD to illustrate the features considered in each kernel. b and c) The local context kernel (*K_LC_*) uses orthographic and shallow linguistic features (POS, lemma, stem) of the tokens located at the left and right (window size of 2) of the candidate entities (EHD3 and MDD). d) The global context kernel (*K_GC_*) is based on the assumption that an association between two entities (in this case EHD3 and MDD) is more likely to be expressed within on of three patterns (fore-between, between, between-after). In this example the association between EHD3 and MDD is expressed in the between pattern. e) In the global context kernel (*K_GC_*) we consider both trigrams and sparse bigrams in each pattern.

**Figure 2.**
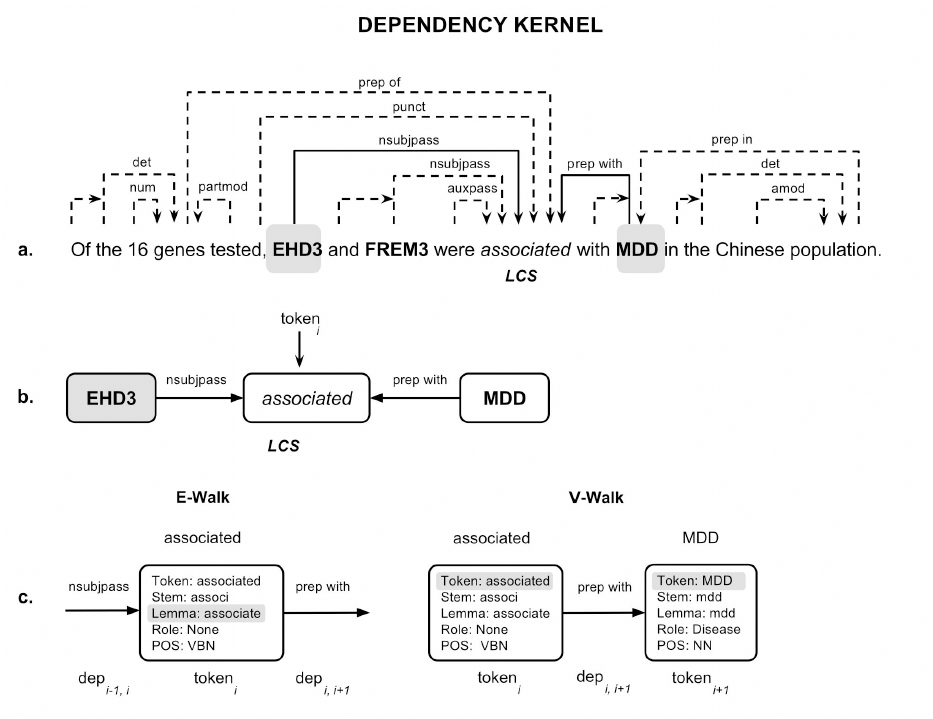
Dependency graph representation of a gene-disease association. a) Dependency graph representation of the sentence. Solid lines represent the shortest path between the two candidates. The token “associated” is the Least Common Subsumer (LCS) node of both candidates. b) Subgraph representing the shortest path between EHD3 and MDD, where syntactic dependencies are represented as edges and tokens as nodes. c) The e-walk and v-walk features for the node “association” and the syntactic (token, stem, lemma, POS) and semantic features (role) considered in the *K_DEP_* kernel.

**Table 1.** Statistics of the EU-ADR and GAD corpora. The Association type classifies the association according to the level of certainty: TRUE (positive (PA), negative (NA) and speculative (SA)) and FALSE (FA).

| Association type |  | EU-ADR |  |  | GAD |
| --- | --- | --- | --- | --- | --- |
|  |  | Drug-Disease | Target-Drug | Gene-Disease |  |
| TRUE | PA | 162 | 157 | 213 | 1833 |
|  | NA | 2 | 6 | 19 | 967 |
|  | SA | 12 | 14 | 30 | - |
| FALSE |  | 68 | 70 | 93 | 2529 |
| TOTAL |  | 244 | 247 | 355 | 5329 |

We then focus on the identification of gene-disease associations. We present the results of the real-life performance of the system and discuss its application for the identification of associations between genes and diseases in two different scenarios: a) the research on the genes involved in depression, one of the major health problems in the world, and b) the population of DisGeNET, a public database of gene-disease associations.

### Identification of drug-target, gene-disease and drug-disease relationships

We assessed the performance of the *K_SL_* and *K_DEP_* kernels on the relationships available in the EU-ADR corpus [22]. We used different combination of features to represent the associations, but only a selection of the better results is shown in the online Suppl. Material (Table 1, http://ibi.imim.es/befree/#supplmaterial) and some of them transcribed here (Table 1). In the case of the drug-disease associations, the best performance both in terms of F-score and Recall is obtained with the *K_DEP_* kernel (3: P 70.2%, R 93.2%, F 79.3%), using stems on the v-walk feature, while in terms of Precision the best result is obtained using POS tags on both the e-walk and v-walk features (19: P 74.5%, R 71.5%, F 72.3%). The best results obtained by combining both kernels did not improve the performance of the dependency kernel alone (75: P 72.0%%, R 84.0%, F 77.0%). Similar results were obtained for the gene-disease association classification, where the *K_DEP_* kernel alone achieved the best performance. The best performance in terms of F-score and Recall was obtained using stem or lemma over the v-walk features (5 and 3: P 75.1 %, R 97.7%, F 84.6%), while the best performance in terms of Precision was obtained when using lemma in v-walk and role in e-walk (30: P 83.8 %, R 71.0%, F 75.6%). Finally, for target-drug relationship, the highest Precision was obtained by the *K_DEP_* kernel with role and POS features over the v-walk and e-walk, respectively (21: P 75.2% R 68.1% F 70.2%), while the highest Recall was obtained when using a combination of *K_SL_* and *K_DEP_* (80: P 73%, R 98% F 82.8%). The best classification in terms of F-score is achieved when using different combination of features with both kernels (see for instance 102: P 74.2%, R 97.4%, F 83.3%). Nevertheless, it is worth to mention that the *K_SL_* kernel, which only uses shallow linguistic information, achieves competitive results in the classification of sentences containing drug-disease, gene-disease and drug-gene associations (F-score: 76.7 %, 80.9%, 81.1% respectively).

The EU-ADR corpus is a valuable resource because it contains annotations for different types of associations, but its main drawback is its small size. There are only a limited number of corpora for entities of biomedical interest (see http://corpora.informatik.hu-berlin.de/ for a recent update). In order to test the feasibility of using a semi-automatic annotated corpus for biomedical relation extraction, we developed a corpus from the GAD database to have a large benchmark of gene-disease associations in which to train and evaluate gene-disease classifiers. Then, we tested the classifier for gene-disease relationships on the GAD corpus by 10-fold cross-validation. Compared to the gene-disease set from the EU-ADR corpus, the GAD corpus is larger and contain a different ratio of true/false associations, thus it is interesting to see how the different combination of kernels and features behave in this benchmark. In addition, it contains a larger fraction of negative sentences, allowing the classification of positive (PA) and negative (NA) sentences pertaining gene-disease associations. Although these annotations are available in the EU-ADR corpus, due to its small size, it did not allow training a classifier to distinguish between positive and negative sentences. When assessing the classification over the class TRUE, the best results where those obtained with the *K_SL_* (1: P 77.8%, R 87.2% F 82.2%). Contrasting with the results obtained on the EU-ADR corpus for gene-disease associations, *K_DEP_* alone did not work very well on the GAD corpus, and the combination of both kernels showed an improvement of the performance but was always lower than the ones obtained with *K_SL_* alone (see http://ibi.imim.es/befree/#supplbefree). In the scenario of the classification over three classes (PA vs NA vs FALSE), although the best performance in terms of Precision or Recall is obtained with combination of kernels, the best F-score is achieved by the *K_SL_* kernel with sparse bigrams, where the Precision and Recall values although not optimal are competitive (2: P 66.0%, R 73.8% F 69.6%). In summary, these results show that a corpus developed by automatic annotation from an expert-curated database on gene-disease associations can produce competitive classifiers, and that the *K_SL_* kernel with shallow linguistic information performs quite well in the classification.

### Evaluation on real-life case studies

#### Case study on genetic basis of depression

Depression is a chronic, recurring, life-threatening disease and the second cause of morbidity worldwide, costing billions of dollars per year to the society [25]. It is currently accepted that a variety of genetic, environmentally-driven epigenetic changes and neurobiological factors play a role in the development of depression; however the exact mechanisms that lead to the disease and affect therapy efficacy are still poorly understood. MEDLINE currently indexes more than 100,000 publications on depression, thus it is a good resource to gather information on genetic determinants of this illness. We performed an evaluation on a real-life setting to test the performance of BeFree to identify genes associated to depression. We evaluated the results in terms of Precision, Recall and F-score of the predictions. Next, we evaluated the quality of extracted information comparing it with what is available in curated resources. We defined a document set of 270 abstracts pertaining to depression that were published during 2012. This document set was processed to identify genes and depression terms with BioNER (see Methods) and the associations between them using gene-disease models trained in EU-ADR and GAD corpora. In both cases, we used the model that in cross-validation achieved the best F-score (for EU-ADR, experiment 3; for GAD, experiment 1 and 2, see http://ibi.imim.es/befree/#supplbefree, Table 2). From a total number of 830 gene-disease associations predicted by the models, we manually reviewed a subset of 100 selected at random to estimate the performance of each model. In the case of the model trained on the EU-ADR, although the Recall was almost perfect (96.6%), we observe a decrease in the Precision of the classification compared to the cross-validation scenario (59.4%). On the other hand, the model trained on the GAD corpus performed better in terms of Precision (70%) but worst in terms of Recall (59.3%) when compared to the cross-validation scenario. A model trained in the GAD corpus is also able to classify sentences containing gene-disease associations as positive, negative and false with F-score of 53.7 % (data not shown). All in all, the model trained in the EU-ADR corpus, despite its small size, performed a better classification of gene-disease sentences in a real case setting (F-score 73.5%). We then carry out a qualitative analysis of the information extracted by BeFree. We compared the genes identified as related to depression using BeFree with the genes already known to be associated to depression available from DisGeNET. The BeFree model trained on the EU-ADR corpus identified 170 genes from the full set of publications, 41 of them available in DisGeNET, whereas the model trained on the GAD corpus retrieved 106 genes, 37 of them were already reported in DisGeNET (Figure 3). More interestingly, the EU-ADR and the GAD models found 129 and 69 genes respectively, not present in DisGeNET, which might represent novel findings that could be introduced in the database. We analysed more deeply the set of genes that were predicted by both methods and were not present in DisGeNET (59 genes) by functional enrichment analysis with GO terms using DAVID [26]. We found significant annotations for terms like synaptic transmission, transmission of nerve impulse, biogenic amine catabolic process, regulation of neurological system process, regulation of cell cycle, regulation of inflammatory response, which are also found for the list of genes from DisGeNET, and are representative of the biology of depression. More interestingly, some of the genes identified by text mining are putatively involved in RNA regulation, RNA splicing and epigenetic regulation, such as *MEG3* (GeneId: 55384), *BDNF antisense RNA* (GeneId: 497258), *DGCR8* (GeneId: 54487), *EXOSC6* (GeneId: 118460), and *GEMIN4* (GeneId: 50628). This is noteworthy since there is an increasing interest in the relationship between the aforementioned processes and the physiopathology of depression. In summary, the application of the BeFree system achieves competitive performance in a real-case scenario and allows the identification of genes related to depression, not previously associated to the disease in specialized databases. More importantly, some of these genes represent novel aspects of the physiopathology of depression.

**Figure 3.**
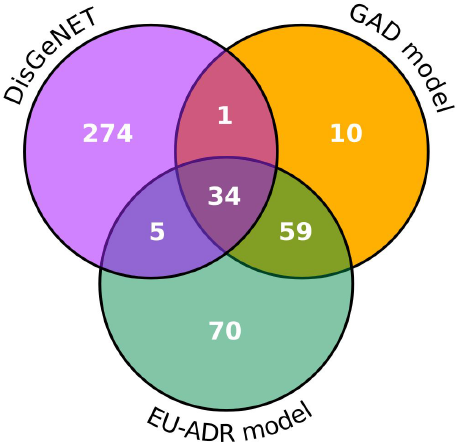
Depression genes identified by BeFree and their overlap with genes available in other repositories. Venn diagram showing the overlap for the depression genes identified by BeFree trained in GAD or EU-ADR corpora, and the depression genes present in DisGeNET.

**Table 2.** Selected results obtained by 10-fold cross-validation on the EU-ADR corpus. The first column indicates the number of the experiment as it appears in http://ibi.imim.es/befree/#supplbefree, Table 1. The second column shows if *K_SL_* is used with (TG+SBG) or without (TG) sparse bigrams, or if it is not used (-). The next two columns focus on *K_DEP_* walk features indicating the use of one of the following features: token (T), stem (S), lemma (L), POS-tag (P), role (R) or none (-). Finally, the last columns show the result obtained in each experiment indicating Precision (P), Recall (R) and f-measure (F) in percentage (%).

| Experiment | $K_{SL}$ | $K_{DEP}$ | | EU-ADR | | | | | | | | |
| --- | --- | --- | --- | --- | --- | --- | --- | --- | --- | --- | --- | --- |
| | | $v$-walk | $e$-walk | Drug-Disease | | | Gene-Disease | | | Target-Drug | | |
|  |  |  |  | P | R | F | P | R | F1 | P | R | F |
| 1 | TG |  |  | 73.4 | 81.0 | 76.7 | 74.2 | 89.6 | 80.9 | 72.7 | 93.6 | 80.8 |
| 2 | TG+SBG |  |  | 72.9 | 79.7 | 75.7 | 73.6 | 89.7 | 80.5 | 72.7 | 94.1 | 81.1 |
| 3 | - | S | - | **70.2** | **93.2** | **79.3** | **75.1** | **97.7** | **84.6** | 72.9 | 97.4 | 82.5 |
| 5 | - | L | - | 70.2 | 92.5 | 79.0 | **75.1** | **97.7** | **84.6** | 72.9 | 97.4 | 82.5 |
| 19 | - | P | P | 74.5 | 71.5 | 72.3 | 74.7 | 53.0 | 59.5 | 72.2 | 67.6 | 68.9 |
| 21 | - | R | P | 72.6 | 70.2 | 70.5 | 77.9 | 53.0 | 61.7 | 75.2 | 68.1 | 70.2 |
| 30 | - | L | R | 66.8 | 65.1 | 63.6 | 83.8 | 71.0 | 75.6 | 71.3 | 73.1 | 70.8 |
| 75 | TG+SBG | L | - | 72.0 | 84.0 | 77.0 | 74.0 | 96.8 | 83.6 | 72.9 | 97.5 | 82.5 |
| 80 | TG+SBG | - | L | 71.9 | 82.1 | 76.3 | 73.8 | 94.8 | 82.7 | 73.0 | 98.0 | 82.8 |
| 102 | TG+SBG | T | R | 72.7 | 81.3 | 76.2 | 75.1 | 91.8 | 82.4 | **74.2** | **97.4** | **83.3** |

#### Large-scale analysis of gene-disease associations from the literature

We applied the BeFree system on a set of 737,712 abstracts pertaining to human diseases and their genes (see Methods for details on document selection) to identify relationships between genes and diseases. Note that our approach for NER takes into account the existing ambiguity in the nomenclature between entities of different semantic type, such as genes and diseases (see Methods for more details). This resulted in 530,347 gene-disease associations between 14,777 genes and 12,650 diseases, which were reported in 355,976 publications. DisGeNET, a database that integrates associations between genes and diseases from several sources, includes 372,465 gene-disease associations at the time of this analysis. Thus, the data identified by BeFree represent a very large dataset on gene-disease associations. Some concerns on the quality of the extracted information could be raised such as 1) errors in the text mining approach, both at the level of NER and RE, and 2) quality of the experimental evidence supporting the association. A simple way to identify both types of error would be to manually curate all the associations, but this is not a feasible task. Thus, before delivering the data to the public through the DisGeNET knowledge portal, we conducted a series of analysis to learn more about the data and its provenance.

##### Data analysis and filtering

We first analysed the frequency distribution of the number of publications or PubMed IDs (PMIDs) that support each disease association (Figure 4). As can be observed from the figure, 68.5% of the associations (363,382 associations) are supported by only one publication and 72,693 by two publications. On the other extreme of the distribution, there are approximately 900 associations supported by more than 200 publications (0,16 %). On average, each association is supported by 2.8 publications. We then inspected in more detail the associations supported by only one publication. These might be associations that could not be reproduced again by any other research group. Alternatively, another reason for the low publication number could be that the related research area is not a hot topic and therefore it is more difficult to publish in this specific domain. On the other hand, these associations could be new findings from recent publications, which might be in the future reproduced or followed up in other publications.

**Figure 4.**
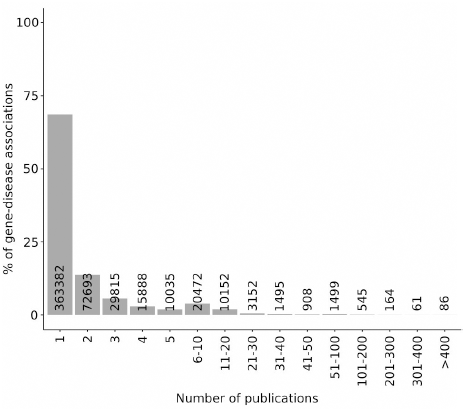
Number of gene-disease associations as a function of the number of PMIDs that support each association.

We analysed in more detail the set of associations supported by only one publication, looking at their publication dates (Figure 5) and the Impact Factors (IF) of the journals (Figure 6), evaluating also their coverage in the DisGeNET database. Figure 5 shows that most of the associations supported by only one publication have been published in the last 15 years. Remarkable, 40,000 associations (11% of all the associations supported by one PMID) have been published during 2013. These associations represent newly open research areas on the genetic basis of diseases that might point out potential candidates for biomarkers or therapeutic targets. Interestingly, almost none (97%) of these associations are present in the DisGeNET database. On the other hand, 35 % of the associations have been published in journals without IF and 16% in journals with IF between 0 and 2.5 (Figure 6). Remarkably, a very small fraction of the associations with one supporting publication have been published in journals with the higher IF, while the majority of the associations are reported in journals with IF between 2.5 and 5.

**Figure 5.**
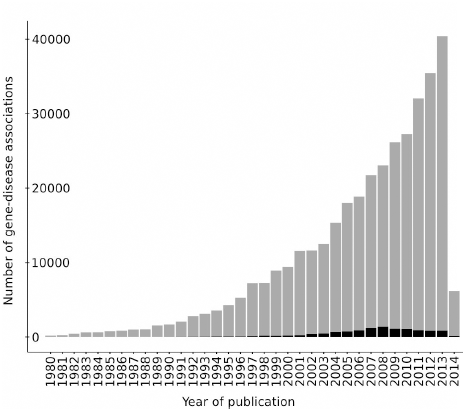
Number of gene-disease associations reported by only one PMID in each calendar year. In red we show the number of associations present in DisGeNET.

**Figure 6.**
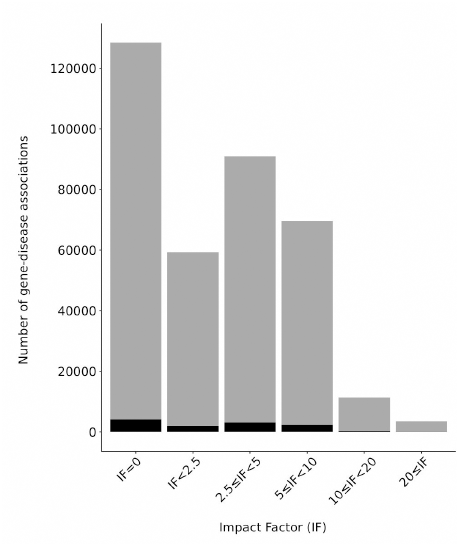
Number of gene-disease associations reported by only one PMID in journals classified by their Impact Factor. In red we show the number of associations present in DisGeNET.

We then inspected the distribution of the number of gene-disease associations reported per MEDLINE abstract (Figure 7). As can be observed, most of the abstracts (47%) report 1-2 gene-disease associations and on average each abstract reports 1.5 gene-disease associations. However, there is a subset of 15 abstracts that report more than 100 gene-disease associations, with one extreme case of 372 gene-disease associations. Manual inspection of the 15 abstracts that report more than 100 associations indicated that in most of the cases they report associations for a number of genes to a disease, using long sentences with coordination structures. In order to avoid possible sources of errors during text mining processing of these long, complex sentences, we decided to remove those abstracts that report more than 20 associations.

**Figure 7.**
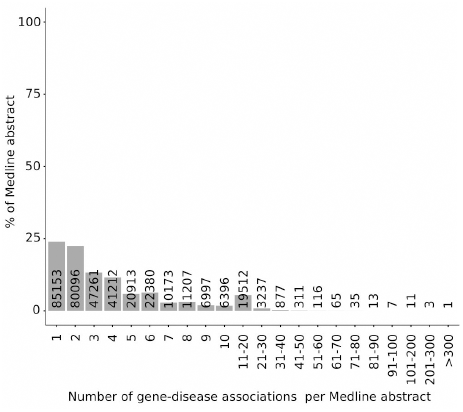
Distribution of the number of gene-disease associations reported per MEDLINE abstract.

Based on this preliminary data analysis, we developed a decision tree workflow on the BeFree data that takes into account the number of publications supporting the gene-disease association, the overlap with DisGeNET and the IF of the journals (Figure 8). After applying this workflow, we obtained 330,888 gene-disease associations (62 % of the original data set) between 13,402 genes and 10,557 diseases.

**Figure 8.**
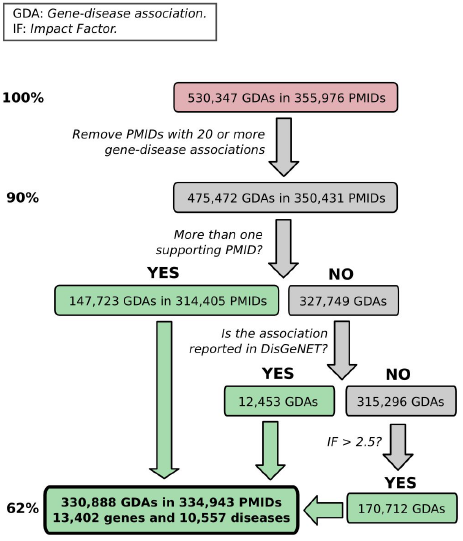
Decision Tree Workflow for selection of BeFree dataset on gene-disease associations.

##### Integration with DisGeNET and data prioritization

A pragmatic way to assess the quality of the extracted information is to contrast it to the information present in expert curated resources. Thus, we integrated the data extracted by Befree with expert reviewed DisGeNET sources (curated and predicted, see http://www.disgenet.org/web/DisGeNET/v2.1/dbinfo#sources for more details on DisGeNET datasets) in order to perform this comparison. Only 7,669 gene-disease associations (2% of BeFree associations) are in common between expert curated associations from DisGeNET and BeFree, while the overlap between the two sources is quite small (0.3% of BeFree associations, Figure 9). Remarkably, from all the gene-disease associations (curated, predicted, and BeFree) 3.9% are only reported by curated sources, and 92.5% only provided by BeFree. The gene-disease associations present in DisGeNET but not recovered by BeFree might be examples of associations mentioned in the full-text and supplementary material of articles and not present in the abstract, or derive from publications not retrieved by our PubMed query used for document selection. Alternatively, they might be false negatives from our text mining approach. The high percentage of associations recovered by text mining and not present in the curated resources highlight the difficulty in collating all this putative useful information in curated databases. It is important to note that in our approach we do not mine the full MEDLINE repository but only a small, but significant in terms of content, fraction (approx. 3 % of current MEDLINE database).

**Figure 9.**
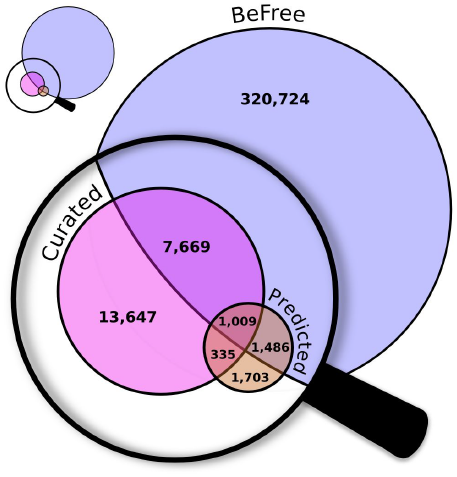
Overlap of the gene-disease associations identified by BeFree with the associations available in DisGeNET curated and predicted sources. DisGeNET information coming from expert curated sources such as UniProt are classified as curated, whereas information coming from model animals such as mouse are classified as predicted. For more information see http://www.disgenet.org/

We computed a score for the gene-disease association based on the number of data sources that report the association, the level of curation of each source and number of supporting publications (see Methods) in order to analyse in an integrative manner the data extracted by text mining. Figure 10 shows the DisGeNET score for the BeFree associations versus the number of supporting publications for each association. Most of the associations (99%) have less than 200 publications and have a wide range of scores, reflecting the fact that they are reported in one or several sources with different levels of curation. Moreover, the analysis of this plot let us identify some interesting outliers. First, the associations *APP*-Alzheimer disease and *CFTR*-Cystic Fibrosis receive a very high score because they are also reported in all the DisGeNET sources, and represent examples of very well studied gene-disease associations. Remarkably, there are 22 associations with very low score (0.06, meaning that they are only reported by BeFree) but are reported by more than 1000 publications. It is intriguing why these associations, that seem to be very well studied ones since are reported in thousands of papers, are not present in any other DisGeNET source. A closer look to some of them (Table 3) indicate that they represent very well studied gene disease associations between breast and ovarian cancer with *TP53*, *BRCA1*, *BRCA2*, *ESR1*, *ERBB2*, and also associations of specific genes to generic cancer terms (neoplasms). For all these cases we find the corresponding gene-disease association in DisGeNET but with a different, yet closely related UMLS concept. The diversity both at expressivity and granularity levels of disease terminologies used in biomedical sources is highlighted by the large number of CUIs normalized (aprox. 10,000) in associations unlocked by text mining. This opens the question about disease terminologies standardization, specially to ensure interoperability in translational research. We observed differences in the disease terminology used by database curators and the literature. In general, there is a preference for using disease concepts that contain MeSH terms by database curators (at least for the databases included in DisGeNET). It is interesting to note that most of the disease concepts present in the curated sources in DisGeNET contain MeSH terms, while this is not the case for the data extracted by BeFree (see Figure 13).

**Figure 10.**
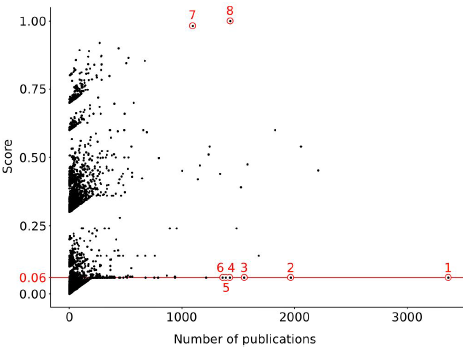
DisGeNET score vs number of supporting publications for the gene-disease associations identified by BeFree. The selected examples discussed in the text are: 1) TP53-Malignant Neoplasm; 2) BRCA1-Breast Carcinoma; 3) ESR1-Breast Carcinoma; 4) ERBB2-Breast Carcinoma; 5) BRCA1-Ovarian Carcinoma; 7) APP-Alzheimer disease; 8) CFTR-Cystic Fibrosis.

**Table 3.**
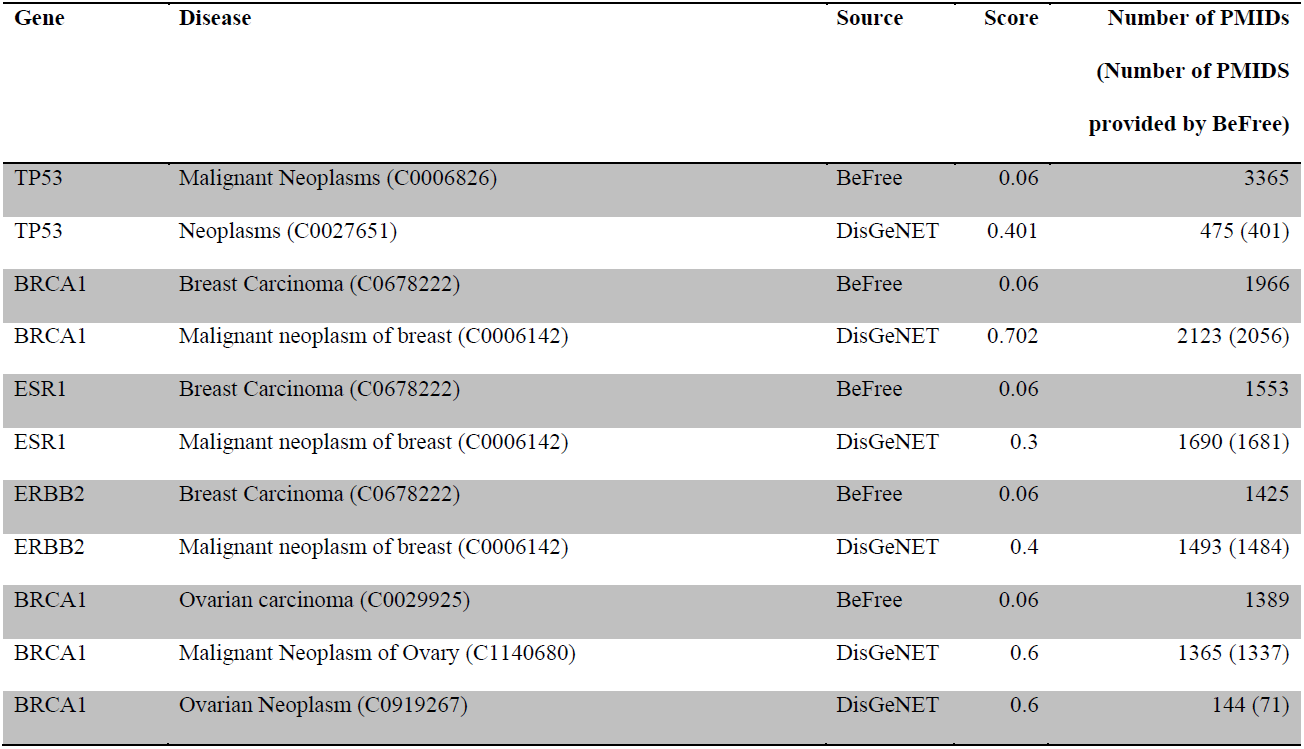
Examples of gene-disease associations from BeFree with low score and supported by a large number of publications. Similar associations provided by DisGeNET are indicated.

##### Characterizatiob of BeFree data

We analysed the frequency distribution of the number of associated diseases per gene (Figure 11) and the number of associated genes per disease (Figure 12). The plots show that there are very “promiscuous” genes regarding their association to diseases (e.g. *VEGFA*, *IL6*, *TNF* and *TP53*), whereas other genes seem to be more specific as they are reported as associated with one or two diseases only. The same consideration can be done when analysing diseases and their associated genes (Figure 12). In this case, it is expected that neoplastic diseases occupy the extremes of the distribution, both for their genetic heterogeneity or for being very well studied. Finally, we inspected the therapeutic areas (Figure 13) and the protein classes covered by the gene-disease associations identified by BeFree (Figure 14). The coverage of diseases by therapeutic areas according to the MeSH classification from BeFree paralleled the one obtained for the diseases in DisGeNET. It is important to note that a large fraction (more than 40%) of diseases identified by BeFree cannot be assigned to a MeSH disease class, while this is not the case for DisGeNET diseases. Thus, the five most covered MeSH disease classes in BeFree, following the not classified diseases, are “Congenital, Hereditary, and Neonatal Diseases Abnormalities”, “Nervous System Diseases”, “Pathological Conditions, Signs and Symptoms”, “Nutritional and Metabolic Diseases and Neoplasms”. Finally, the disease genes identified by BeFree are classified as protein-coding (89%), ncRNA (2%), pseudogenes (2%), rRNA, tRNA, snoRNA and snRNA (0.5%), other (6%). Again, regarding the classification of the proteins encoded by disease genes according to the Panther protein classification, we observe again that disease proteins identified by BeFree have a similar class distribution than those present in DisGeNET (Figure 14).

**Figure 11.**
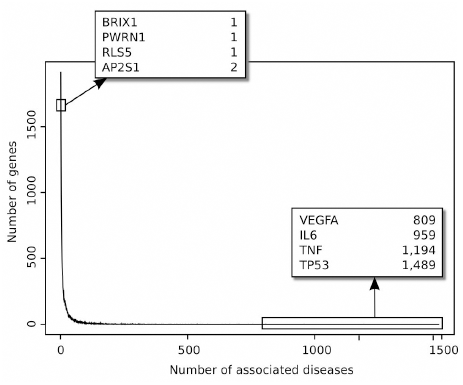
Frequency distribution of the number of associated diseases per gene.

**Figure 12.**
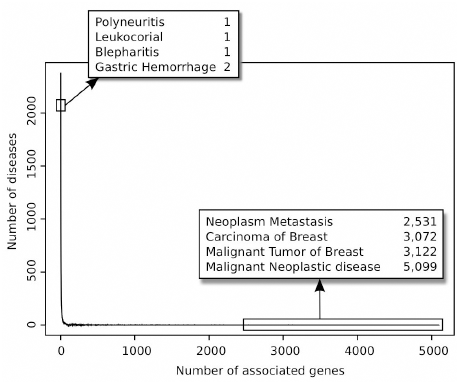
Frequency distribution of the number of associated genes per disease.

**Figure 13.**
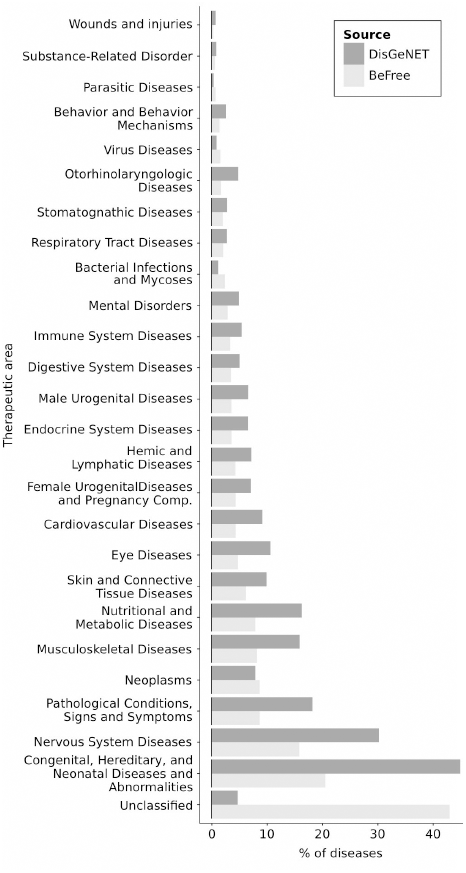
Distribution of diseases according to the MeSH disease classification in the BeFree and DisGeNET datasets. Note that more than 40% of diseases in BeFree do not contain a MeSH disease class.

**Figure 14.**
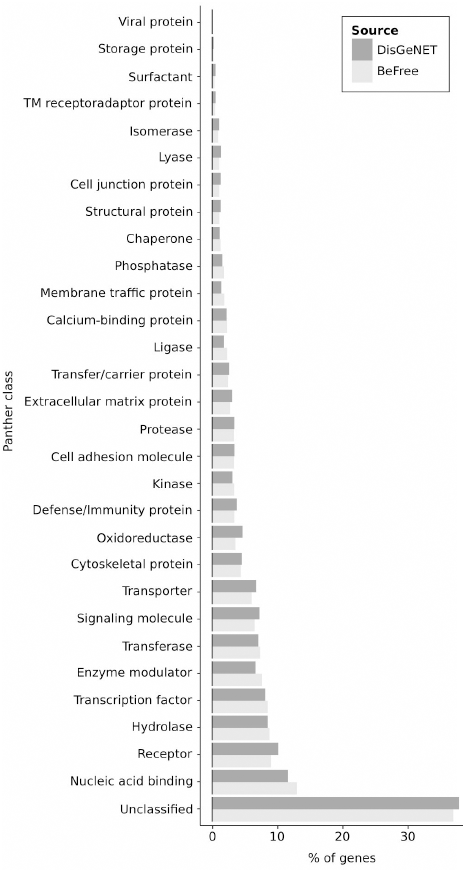
Distribution of disease proteins according to the Panther Protein classification. Data from Panther (http://www.pantherdb.org/) was used to annotate disease proteins from BeFree and DisGeNET. Note that more than 37% of proteins in BeFree cannot be classified according to Panther.

## Conclusions

Our results show that a kernel based approach using both morpho-syntactic and dependency information performs competitively for the identification of drug-disease, drug-target and gene-disease relationships from free text. Although the exact combination of features that yield better results depends both on the association type and the corpus used for training the system, the use of shallow linguistic information is enough to produce accurate RE classifiers to recognize these associations. A RE system for gene-disease associations trained on different corpora (EU-ADR, GAD) with very different characteristics is able to identify gene-disease associations in real-case scenarios with good performance. Finally, as previously suggested by others [5], a corpus developed by semi-automatic annotation is a good resource for developing a RE system in biomedicine.

In addition, we evaluated the value of the information extracted by BeFree for specific case studies in translational research. Particularly, the results obtained in the case study on depression indicated that BeFree is able to identify genes associated to depression that are not present in public databases and support novel hypothesis in the pathophysiology of depression. The large-scale analysis of gene-disease associations provided interesting insights on the kind of information that can be found in the literature about gene-disease associations, and raised some issues regarding data prioritization and curation. The conclusions of the analysis of the provenance of the gene-disease associations identified by BeFree can be summarized in the following points:

- The scientific literature is a rich resource for extracting gene-disease associations, even considering only abstracts from a specific subset of MEDLINE.
- Only a small proportion of the gene-disease associations discovered by text mining are collected in expert-curated databases. There is a pressing need to find alternative strategies to manual curation to review, prioritize and curate these associations and incorporate them into domain-specific databases.
- A first and important step is to extract this information and put it in a standardized format to allow its integration with other data sources and their subsequent analysis for different purposes.
- Joint analysis of data derived by text mining with data curated by experts appears as a suitable approach to assess data quality and identify novel and interesting information.
- A large proportion of the associations are supported by only one publication, raising concerns on data reproducibility but also pointing out novel putative targets for research and innovation.
- There are important differences in the use of disease terminologies between database curators and the authors of publications, and also in the level of granularity of disease concepts to describe a disease phenotype. This is a current challenge for large-scale disease data integration that aims to gather a comprehensive coverage of disease and ensure systematic interoperability across biomedical domains.

Biocuration of large data sets, a.k.a. big data, is becoming a bottleneck for biomedical research. Recently, the crowdsourcing approach has attracted interest in the bioinformatic domain and holds promise for biocuration tasks [27]. As more and more scientific groups are extracting knowledge blocked in free text by text mining and exposing it to the public domain, another upcoming question is the meta-curation of such deluge of data. In this regard, the nanopublication concept, based on semantic web triple-assertions, is a promising approach to aid the prioritization of associations based on the supporting evidence [28]. These kind of approaches could be applied to large datasets such as the gene-disease associations extracted by BeFree.

In summary, the study presented here highlight the importance of performing several steps of data analysis on large data sets, before using the data for further bioinformatic analysis and even to feed it to the curation pipeline of a database. We suggest that this kind of iterative process of data extraction, analysis and refinement of data extraction methodology should be applied to other approaches aimed at extracting large-scale information from the literature.

## Methods

We present the development of the BeFree system, composed of a biomedical named entity recognition system (BioNER, presented in [23] and a kernel-based relation extraction module that is described below.

### Kernel based Relation Extraction

In order to implement a RE for different relationships (drug-target, drug-disease, gene-disease), we propose the combination of the Shallow Linguistic Kernel (*K_SL_*) based on the system originally proposed by [17] and our Dependency Kernel (*K_DEP_*). Both kernels are described in the next two sections.

### Shallow Linguistic Kernel (*K_SL_*)

The Shallow Linguistic Kernel (*K_SL_*), developed by [17] has been successfully applied to PPI, drug-side effects [9] and drug-drug interaction extraction [8]. Here, we propose its application for the identification of the relationships gene-disease, drug-disease and drug-target. *K_SL_* is composed of a linear combination of the kernels *K_GC_* and *K_LC_* that provide different representations of the association between two candidate entities. The global context kernel (*K_GC_*) is based on the assumption that an association between two entities is more likely to be expressed within one of three patterns (fore-between, between, between-after, see Figure 1.d). Three term frequency vectors are obtained based on the bag-of-words approach using trigrams of tokens. Sparse bigrams were included to improve the classification performance, as suggested in the original implementation. The local context kernel (*K_LC_*) uses orthographic and shallow linguistic features (POS, lemma, stem) of the tokens located at the left and right of the candidate entities (window size of 2). Figure 1 shows the features considered by each kernel using an example sentence.

### Dependency Kernel (*K_DEP_*)

We developed the Dependency Kernel (*K_DEP_*) to train a model to recognize relationships between the entities of interest using walk features [20]. The syntactic dependencies of the words within a sentence can be represented as dependency graphs. Figure 2.a shows the dependency graph of an example sentence extracted from MEDLINE as obtained by the Stanford parser (http://nlp.stanford.edu/software/lex-parser.shtml). The shortest path between the two candidate entities can be extracted from the dependency graph (highlighted with a solid line in the example, Fig.2b), which includes the Least Common Subsumer (LCS) node (common governor node between the two candidates, subgraph detailed in Figure 2.b). In the *K_DEP_*, two types of walk features are used, the v-walk feature that is composed of *node_(i)_-edge_(i, i+1)_-node_(i+1)_,* and the e-walk feature that is composed of *edge_(i-1, i)_-node_(i)_-edge_(i, i+1)_* (both illustrated in Figure 2.c). For the edges we consider the dependency relation type, while for the nodes we consider different features of the token, such as the token itself, its stem, lemma, role (if this token is candidate or not) and part-of-speech (POS) tag.

### Corpora

We used two manually annotated corpora: AIMed for PPIs and EU-ADR for gene-disease, drug-target and disease-drug associations. In addition, we developed a semi-automatically annotated corpus for gene-disease associations based on the GAD database. All datasets were pre-processed with a combination of tools to extract the features required by the relation extraction system. More specifically, sentence boundaries were identified by NLTK (http://www.nltk.org/), tokens and part-of-speech (POS) tags were obtained using UIMA modules (http://www.julielab.de/), lemmas were obtained with Biolemmatizer (http://biolemmatizer.sourceforge.net/) and stems were identified with the Porter's algorithm. Finally, syntactic dependencies were obtained with the Stanford parser (http://nlp.stanford.edu/software/lex-parser.shtml). Both EU-ADR and GAD corpora are publicly available (http://ibi.imim.es/befree/#corpora).

### AIMed

The AIMed corpus is widely used for PPI extraction (ftp://ftp.cs.utexas.edu/pub/mooney/bio-data/). The AImed corpus consists of 225 MEDLINE abstracts, of which 200 abstracts describe interactions between human proteins and 25 do not refer to any interaction. There are 5625 annotated sentences, 1008 containing a true PPI (TRUE) and 4617 not containing a true PPI (FALSE).

### EU-ADR

The EU-ADR corpus contains annotations of different entities (drugs, diseases, and genes/proteins) and the relationships between them [22]. In particular, it contains annotations of relationships between drug and diseases (drug-disease set), drug and their protein targets (drug-target set) and genes/proteins and their association to diseases (gene-disease set). In addition, each relationship is classified according to its level of certainty as: positive association (PA), negative association (NA), speculative association (SA) and false association (FA). The EU-ADR corpus is composed of 100 MEDLINE abstracts for each relationship set, and its annotation was performed by three experts. In this study we considered the relationships that result from the consensus annotation of two experts. Table 1 shows the number of relationships for each set.

### GAD

The Genetic Association Database (GAD) is an archive of human genetic association studies of complex diseases, including summary data extracted from publications on candidate gene and GWAS studies (http://geneticassociationdb.nih.gov/). We use GAD for the development of a corpus on associations between genes and diseases (downloaded on January 21st, 2013). We considered the annotations of relationships between a gene and a disease in a single sentence as a reference set to build this corpus. GAD contains over 130,000 records with different type of information. We selected the records satisfying the following requirements: (i) the association between gene and disease is annotated as positive or negative, (ii) the association is expressed in one sentence and (iii) the Entrez Gene identifier for the gene is provided. Although GAD provides the sentence in which a gene-disease association is stated, there is no information on the exact location of the gene and disease entities in the text. In order to develop a corpus suitable for training a gene-disease relation extraction system, the exact location of the interacting entities in the text is required. To achieve that, we applied our own NER system (BioNER, see below) to identify the gene and disease entities in the text and normalize them to NCBI Gene and UMLS identifiers, respectively. Then, the sentences in which a given gene was found together with a specific disease, and this gene-disease association was annotated by GAD curators as positive or negative were labelled as TRUE. In order to create a dataset containing false associations (FALSE) between a gene and a disease, that is, a gene and a disease that co-occur in a sentence but are semantically not associated, we selected the sentences with co-occurrences between a disease and a gene found by the BioNER system that were not annotated by GAD curators as gene-disease associations. Table S2 shows the number of TRUE and FALSE associations that represent the GAD corpus.

### Evaluation of Kernel based Relation Extraction

The performance of each model for association classification was evaluated by sentence-level 10-fold cross validation in each corpus. The classifiers’ performances were assessed using P, R and F-score over the class TRUE. TRUE sentences contain real relationship between the entities analysed, in contrast with FALSE sentences where the two entities co-occur, but there is no semantic relationship between them. In the case of the GAD corpus, we also trained a classifier that distinguishes between positive, negative and false associations, and therefore the performance was assessed over the class positive (PA) and negative (NA) separately.

### Identification of entities

We identified gene and disease mentions in free text using the BioNER system [23]. Briefly, BioNER uses gene and disease dictionaries with fuzzy and pattern matching methods to find and uniquely identify these entity mentions in the literature. During initial analysis of the RE results we observed that a source of error was the wrong identification of entities due to ambiguities in the terminologies for diseases and genes. This is particularly problematic in the case of acronyms, where the same token can be used to refer to a disease or a gene. Thus, we introduced a series of modifications on BioNER in order to address ambiguities in the identification of a single entity type (e.g a gene) and between different entity types (genes and diseases).

Frequently, an acronym appears after the long term is defined in the text. In this case, we compare the list of concept identifiers of both mentions (acronym, long form) to figure out if the acronym refers to the long form. For example, in the sentence “Selective gene targeting using the carcinoembryonic antigen (CEA) promoter is useful in gene therapy for gastrointestinal cancer” (from PMID 11053994), BioNER detects the long form expression “carcinoembryonic antigen” as a gene with NCBI Gene Id 1048, and the acronym “CEA” as four different gene entities (with four NCBI Gene Ids 1087, 5670, 1084 and 1048). The concept identifier in common between the two entities (NCBI Gene Id 1048) is kept as the right annotation. If there is more than one concept identifier in common between the two entities, we look at the similarity of the terms of each concept to select the right identifier. The file gene2pubmed source from Entrez Gene was also used to select the correct identifier in these ambiguous cases. Evaluation of BioNER for gene normalization using the BioCreative II Gene Normalization (BC2GN) [29] resulted in very low Precision (P: 48.1% R: 80.1% F: 60.1%), which could be improved considerably when applying the above mentioned strategies to handle the ambiguities between genes (P: 74.0%, R: 76.2%, F: 75.0%).

The other type of ambiguity arises when one candidate entity in a sentence can refer to different semantic types (disease and gene). For example the symbol “APC” can refer to the gene “adenomatous polyposis coli” or to the disease “atrial premature complex”. To properly recognize the identity of the mention, we take into account the contextual information of the candidate entity. For instance, to disambiguate a candidate entity to a gene, we look for keywords such as “gene”, “protein”, “factor”, “target”, “biomarker”, etc., whereas to disambiguate a candidate entity to a disease, we look for keywords like “disease”, “disorder”, “condition”, “syndrome”, etc. Finally, we also looked at the MeSH Disease annotations of the corresponding abstract to decide if a candidate entity refers to a gene or a disease. We compared the terms of the candidate entity to the terms of the MeSH disease concepts annotated to the abstract using a soft-matching approach, and if a match was found, we annotated the candidate entity as a disease.

In addition, we also performed an evaluation of the performance of BioNER in the identification and normalization of disease entities using the Arizona Disease Corpus achieving competitive (P:72.1% R:64.4% F:68.0%) results compared to previous approaches [30, 31].

### Case study on genetic basis of depression

We defined a PubMed query to retrieve a set of document to depression and published in 2012 as follows:

> (“Depression”[Mesh] OR “Depressive Disorder”[Mesh]) AND “genetics”[Subheading] AND (hasabstract[text] AND (“2012”[PDAT]) AND English[lang] AND “humans”[MeSH Terms]) NOT (“Case Reports”[PT] OR “Clinical Trial”[PT] OR “Clinical conference”[PT] OR “Clinical Trial, Phase I”[PT] OR “Clinical Trial, Phase II”[PT] OR “Clinical Trial, Phase III”[PT] OR “Clinical Trial, Phase IV”[PT] OR “Controlled Clinical Trial”[PT] OR “Randomized Controlled Trial”[PT] OR “Meta-Analysis”[PT])

This query resulted in 270 citations (date of search March 19, 2013). The abstracts were processed with BeFree trained on GAD and EU-ADR corpora to find gene-disease associations.

### Case study on large-scale analysis of gene-disease associations from the literature

We defined a PubMed query to retrieve documents pertaining to human diseases and their associated genes published from 1980:

> (“Psychiatry and Psychology Category”[Mesh] AND “genetics”[Subheading]) OR (“Diseases Category”[Mesh] AND “genetics”[Subheading]) AND (hasabstract[text] AND (“1980”[PDAT]: “2014”[PDAT]) AND “humans”[MeSH Terms] AND English[lang])

This query retrieved 737,712 citations, which were processed by Befree trained on the EU-ADR corpus to identify relationships between genes and diseases.

### DisGeNET score

The DisGeNET score is described in detail elsewhere (Piñero et al, DisGeNET: a platform for the dynamical exploration of human diseases and their genes, *submitted*). Here we reproduce its formulation in order to help in the interpretation of the results. Briefly, we assign a score to each gene-disease association in DisGeNET [24] according to the source in which this association is reported (CURATED, PREDICTED, LITERATURE), the level of curation of each source, and the number of publications that report each association in the case of LITERATURE sources. DisGeNET is a database on gene-disease associations covering all therapeutic areas, that integrates information from resources curated by human experts (UniProt and CTD), from orthologous genes from mouse and rat (MGD, RGD and CTD), and from literature repositories by text mining (LHGDN and GAD). Thus, the DisGeNET source databases are classified accordingly in CURATED, PREDICTED and LITERATURE reflecting the different sources where each association is reported. The gene-disease associations extracted by BeFree are then classified as LITERATURE once integrated in DisGeNET. For the associations reported in LITERATURE sources, we can rank the associations based on the number of publications that support each association. The DisGeNET score is defined as follows:

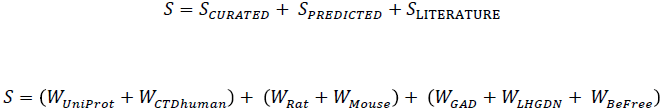

Where

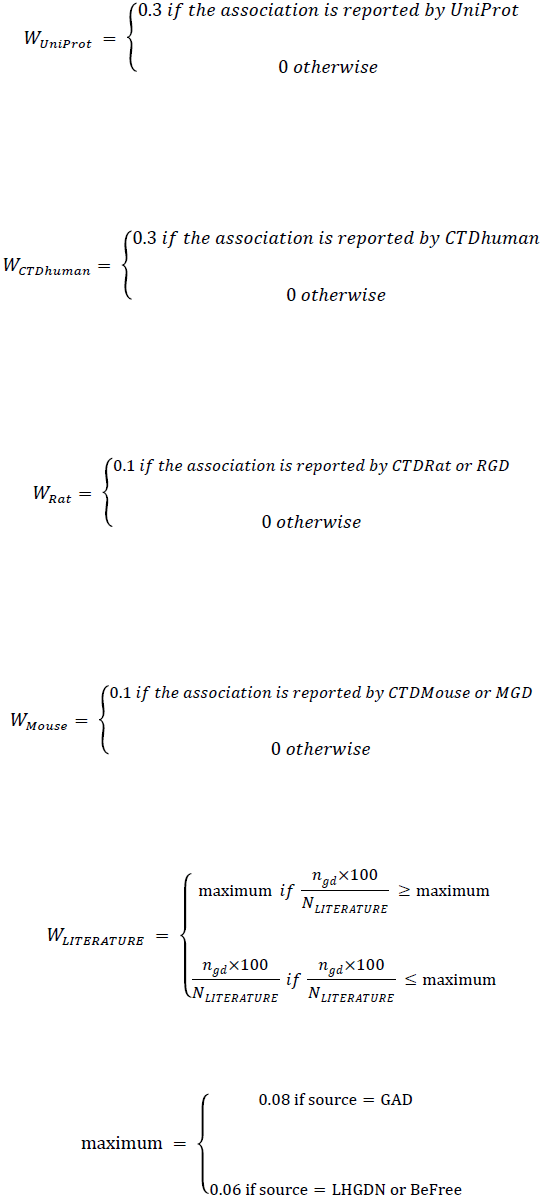

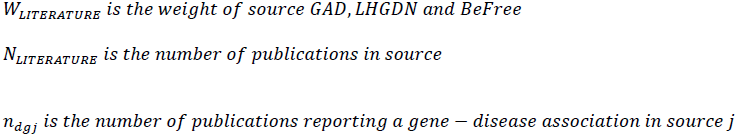

For more details on the DisGeNET score visit the DisGeNET web page (http://www.disgenet.org/).

## Availability

The complete set of gene-disease associations extracted by BeFree, with the supporting statements and information on the provenance, are available in DisGeNET (http://www.disgenet.org). The corpora used in this study are available at http://ibi.imim.es/befree/#corpora.

## Authors' contributions

AB developed the system and performed the evaluation, and participated in data analysis and writing the manuscript. MR contributed to the development of the system. NQ and JP performed data analysis and participated in writing the manuscript. LIF conceived the study, coordinated the work, contributed to data analysis and wrote the manuscript. All authors read and approved the final manuscript.

## Acknowledgements

We thank Dr. Olga Valverde (UPF) and Dr. Marta Torrens (IMIM) for their expertise in the use case of depression. Funding: The research leading to these results has received support from Instituto de Salud Carlos III-Fondo Europeo de Desarollo Regional (PI13/00082 and PI13/00082), the Innovative Medicines Initiative Joint Undertaking under grants agreements n° [115002] (eTOX) and n° [115191] (Open PHACTS)], resources of which are composed of financial contribution from the European Union's Seventh Framework Programme (FP7/2007-2013) and EFPIA companies’ in kind contribution. À.B. and L.I.F received support from Instituto de Salud Carlos III Fondo Europeo de Desarollo Regional (CP10/00524). The Research Programme on Biomedical Informatics (GRIB) is a node of the Spanish National Institute of Bioinformatics (INB).

## Additional files

**Additional file S1 – Evaluation of the RE system on the AIMED corpus**

In addition, this article contains supplementary information available online (http://ibi.imim.es/befree/#supplmaterial).

